# Human PRPS1 filaments stabilize allosteric sites to regulate activity

**DOI:** 10.1101/2022.06.27.496812

**Authors:** Kelli L. Hvorecny, Kenzee A. Hargett, Joel Quispe, Justin M. Kollman

## Abstract

The universally conserved enzyme phosphoribosyl pyrophosphate synthetase (PRPS) assembles filaments in evolutionarily diverse organisms. PRPS is a key regulator of nucleotide metabolism, and mutations in the human enzyme PRPS1 lead to a spectrum of diseases. Here, we determine structures of PRPS1 filaments in active and inhibited states, with fixed assembly contacts accommodating both conformations. The conserved assembly interface stabilizes the binding site for the essential activator phosphate, increasing activity in the filament. Some disease mutations alter assembly, supporting the link between filament stability and activity. Structures of active PRPS1 filaments turning over substrate also reveal coupling of catalysis in one active site with product release in an adjacent site. PRPS1 filaments therefore provide an additional layer of allosteric control, conserved throughout evolution, with likely impact on metabolic homeostasis. Stabilization of allosteric binding sites by polymerization adds to the growing diversity of assembly-based enzyme regulatory mechanisms.

## Introduction

Phosphoribosyl pyrophosphate synthetase (PRPS) catalyzes the production of phosphoribosyl pyrophosphate (PRPP), which in humans is required primarily for nucleotide biosynthesis. The highly conserved enzyme is found in organisms across all the domains of life^1,2^. Humans have three copies of PRPS, with PRPS1 expressed in most human tissues^3^. Mutations in PRPS1 lead to a spectrum of diseases, with gain-of-function mutations producing excess levels of uric acid that cause hyperuricemia and gout, and patients with loss-of-function mutations deficient in nucleotide production with a range of neurological phenotypes, which include deafness, Charcot-Marie-Tooth disorder, and Arts syndrome^4–6^.

PRPS is a critical regulatory node linking pentose phosphate pathway to nucleotide biosynthesis. The enzyme transfers pyrophosphate from ATP to ribose-5-phosphate (R5P), a product of the pentose phosphate pathway, producing PRPP and AMP. Magnesium is required as a cofactor for the reaction, and phosphate is a required allosteric activator (Figure 1A)^2,7,8^. PRPP is essential for *de novo* purine and pyrimidine nucleotide biosynthesis pathways, and for nucleotide salvage pathways. Given the central role of PRPP in maintaining nucleotide levels, its production by PRPS is tightly controlled at multiple levels. Several downstream products inhibit PRPS activity, including the allosteric inhibitors ADP and GDP^9^.

**Figure 1.**
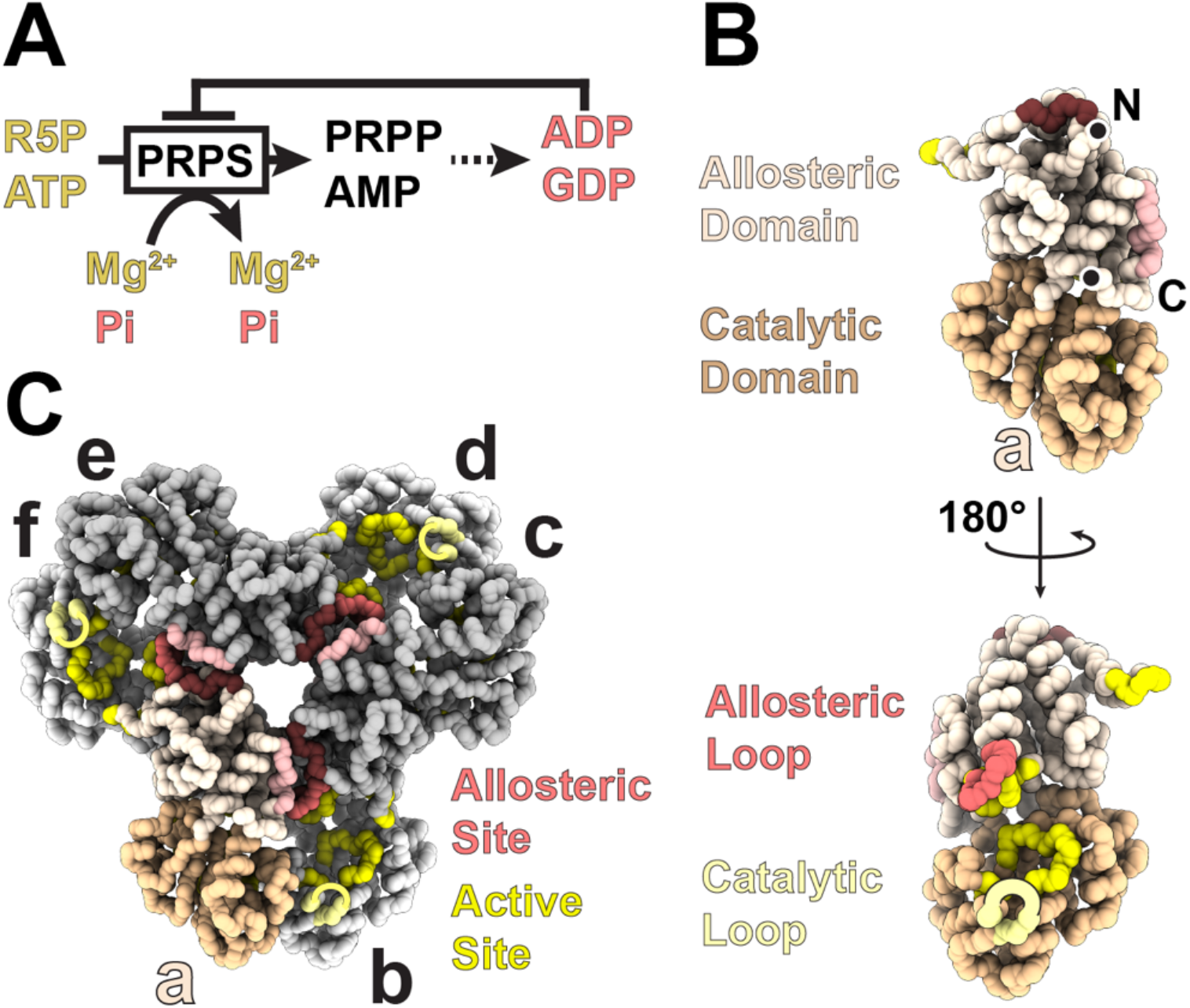
Biochemical and Structural Overview of Phosphoribosyl Pyrophosphate Synthetase. A. Schematic of PRPS catalysis and regulation. B. Backbone trace of a PRPS1 protomer, showing domain organization and residues contributing to allosteric and catalytic sites. C. PRPS1 hexamer, showing how multiple chains contribute to each allosteric and active site.

PRPS assembles into hexamers, which enables formation of both the active site and the allosteric binding site for ADP and phosphate^10,11^. PRPS protomers each have an allosteric domain (containing the allosteric loop) and a catalytic domain (containing the catalytic loop) (Figure 1B). Protomers assemble via their allosteric domains to produce a “bowed” dimer (the b-c dimer in Figure 1C) in solution, and in doing so create the catalytic site where ATP, ribose-5-phosphate, and magnesium bind. Three of these dimers assemble to form the hexamer, creating the allosteric site with residues from three different protomers (Figure 1C). This single site binds both phosphate (activator) or ADP (inhibitor).

Foundational papers looking at PRPS from human tissue document the enzyme’s ability to form large, reversible “aggregates”^8,12^. The large oligomers were observed in activating conditions in the presence of phosphate and Mg-ATP, as well as in conditions that included the inhibitor ADP^13^. The “aggregates” have been long thought to be the active enzyme. Several studies applied a combination of size exclusion chromatography and analytical ultracentrifugation to quantify the number of subunits in the large, active form of the enzyme, suggesting that assemblies larger than a single hexamer were the active form^13,14^. Others linked lower aggregate formation with disease phenotype^15^. Subsequent crystal structures of PRPS hexamers from several different organisms provided significant insight into catalytic and regulatory mechanisms, and the functional role of larger oligomers has been mostly neglected in recent years^10,11,16–18^. More recent work *in vivo* shows that PRPS assembles into micron-scale filamentous polymers in evolutionarily diverse organisms, including humans, *Drosophila*, budding yeast, and *E. coli*, suggesting that these higher order structures play a crucial and conserved role in the cell^19–21^. These new cellular observations have revived interest in understanding the functional role of higher order PRPS assembly, especially given recent progress in understanding the regulatory role of filament assembly in multiple other metabolic enzymes^22–24^.

Here, we characterize filament formation by human PRPS1, reconstituting filament assembly *in vitro*, and showing that allosteric ligands strongly promote assembly. Cryo-EM structures of the filaments reveal a highly conserved assembly interface that mediates interactions between stacked PRPS1 hexamers. Engineered mutations at this interface severely reduce enzyme activity, suggesting that the role of filaments is to increase PRPP production. Comparison of assembled and unassembled PRPS1 structures shows that filament assembly stabilizes the allosteric site, promoting binding of the essential activator phosphate and increased activity. Loss-of-function disease mutations near the filament interface also disrupt assembly, likely explaining their reduced activity, and suggesting that the ability of PRPS1 to assemble into filaments plays a critical role in human health. Finally, cryo-EM structures of PRPS1 filaments actively turning over substrates provide insight into catalytic mechanisms, including a mechanism for “reloading” the active site through coupled opening and closing in adjacent protomers.

## Results

### Allosteric ligands promote PRPS1 filament formation

Phosphate is strictly required for PRPS activity, and early studies showed that it induces formation of “larger aggregates” of the enzyme^25–27^. Consistent with these earlier studies, we found that purifying recombinant PRPS1 in a phosphate-free buffer yields mostly dimers, and addition of phosphate creates a mixed population of species, including short linear polymers, seen in negative stain EM and size exclusion chromatography (Fig. 2A, Ext. Data Fig. 1). Addition of the substrates ATP or ribose-5-phosphate had little effect on the distribution of oligomeric states, but addition of the allosteric inhibitors ADP or GDP appeared to stabilize longer filaments (Fig. 2A, Ext. Data Fig. 1B).

**Figure 2.**
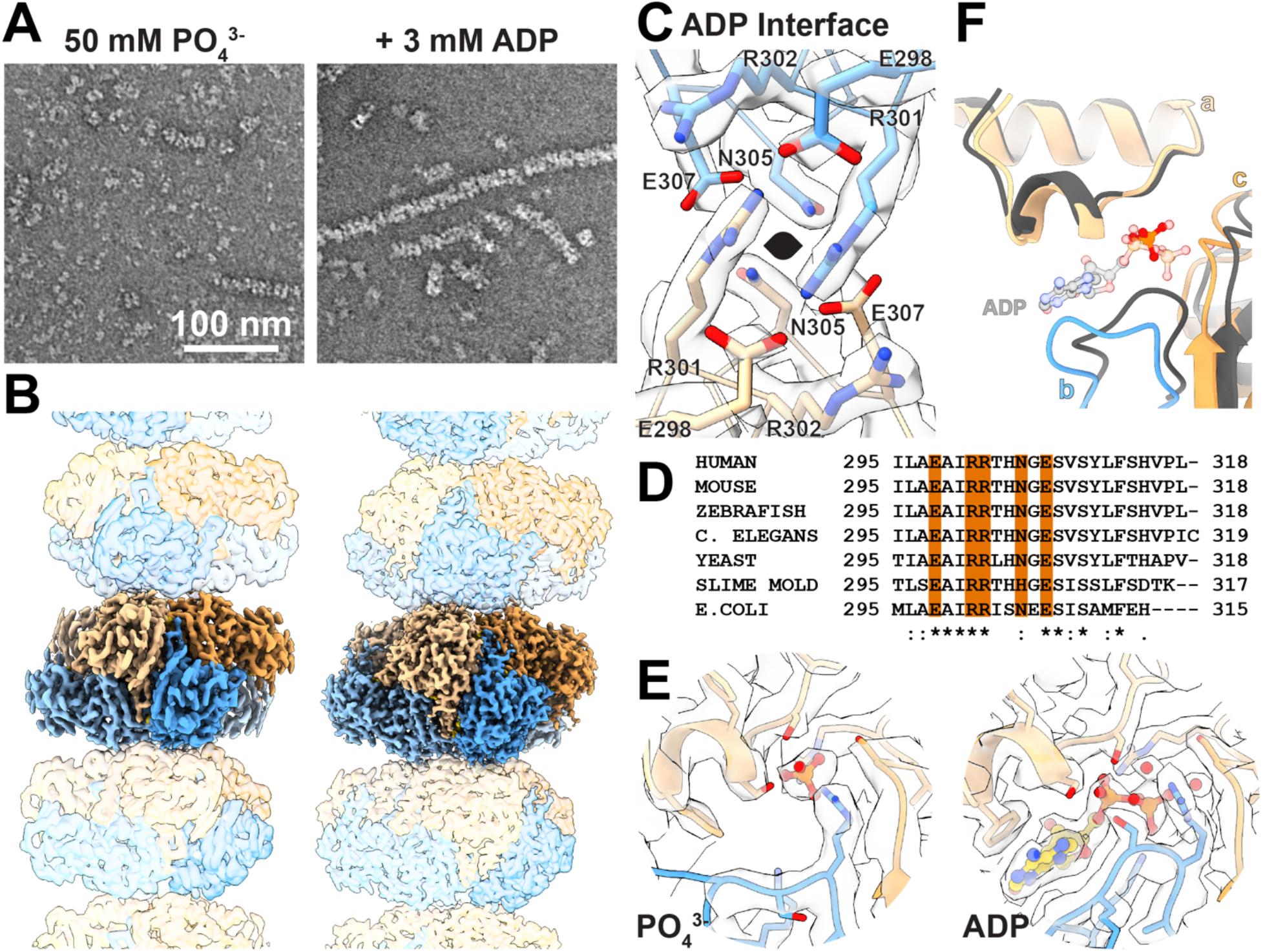
Presence of Phosphate or ADP dictate filament structure of PRPS1. A. Negative stain electron micrographs showing PRPS1 in the presence of phosphate (pH 7.6) or phosphate (pH 7.6) and ADP. B. Cryo-EM structure of PRPS1 filaments bound to phosphate (left) or ADP (right); protomers colored in blue or orange. C. Model and map of the primary interface residues in the ADP-bound structure, with two-fold symmetry axis indicated. D. Sequence alignment showing conservation of the amino acids at the filament interface. E. Model and map of the allosteric sites in the phosphate- and ADP-bound filaments. F. Ribbon diagram and ligands in the allosteric site show that when aligned on the allosteric domain of protomer a, the ligand present dictates the positioning of protomers b and c (phosphate-bound in orange/blue; ADP-bound in grey).

To understand the basis for PRPS1 filament assembly, we determined cryo-EM structures of the phosphate- and ADP-bound enzyme to 3.2 Å and 2.1 Å resolution, respectively (Fig.2B, Ext. Data Fig. 2, Table 1). Both filaments are helices of stacked hexamers with the three-fold symmetry axis coincident with the helical symmetry axis (Figure 2B). Our reconstruction strategy used a single-particle approach, which allowed us to reconstruct short segments of filaments treating them as single particles, followed by local refinement centered either on single hexamers or centered on the filament assembly interface (Ext. Data Fig 2). This approach allows us to determine the highest resolution reconstructions for areas of interest, and has proven useful in other reconstructions of helical filaments^28–30^.

**Data Table 1:**
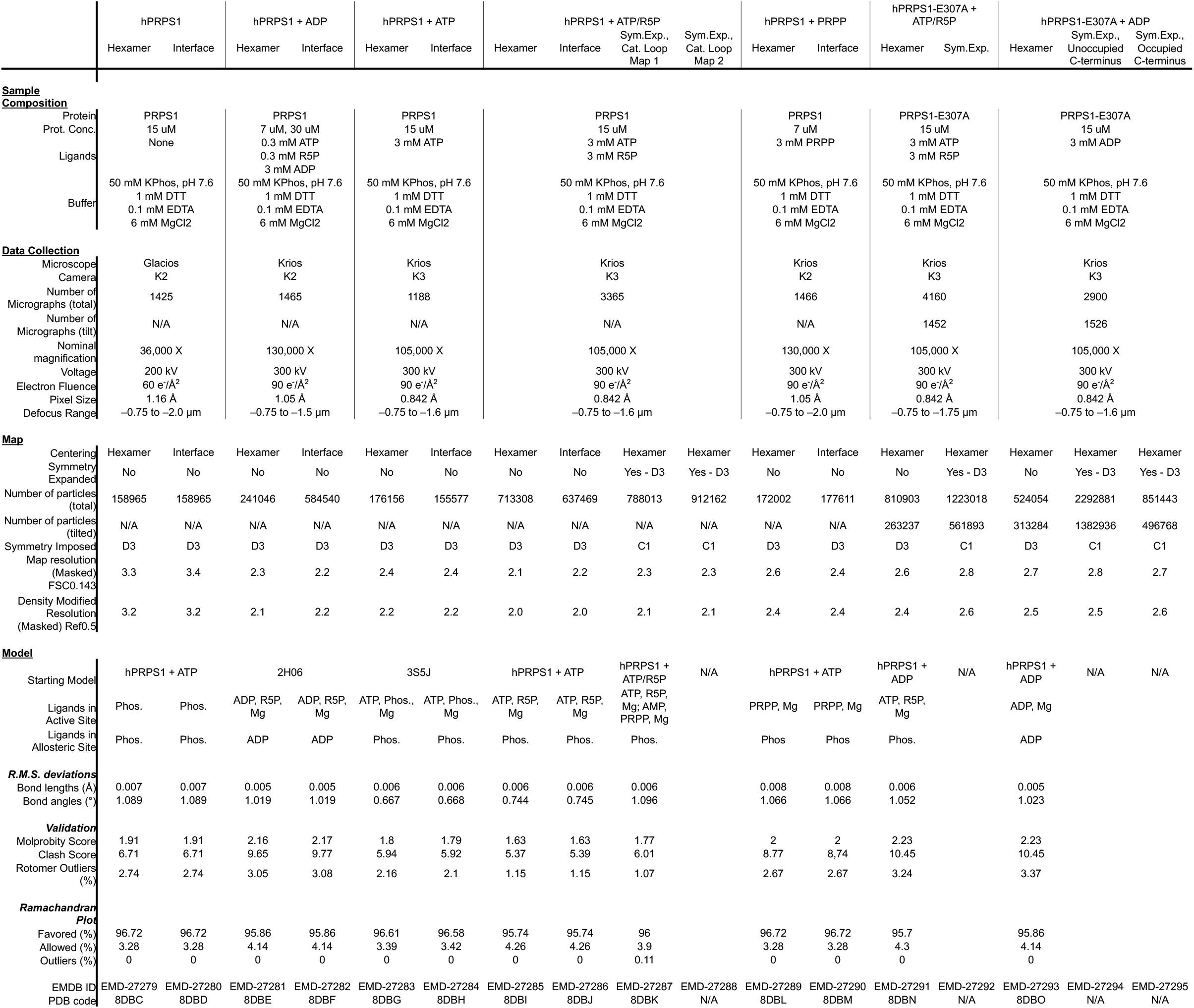
Summary, statistics, and database codes for all volumes and models.

The filament assembly interface is nearly identical in the ADP- and phosphate-bound structures. The primary interface consists of residues E298, R301, R302, N305, and E307, which create a complex network of hydrogen bonds and pi-pi interactions between arginines with the same set of residues in the neighboring hexamer (Figure 2C, Ext. Data Fig. 3A-C). All of these residues are deeply conserved throughout eukaryotes and bacteria (Figure 2D). An additional symmetric contact across the interface is made by N3, with either asparagine or aspartic acid occupying a similar position in other species. Each interaction buries approximately 500 Å^2^ of surface area, and the interactions are repeated three times around the 3-fold symmetry axis, for a total of 1500 Å^2^ between two hexamers in the filament.

**Figure 3.**
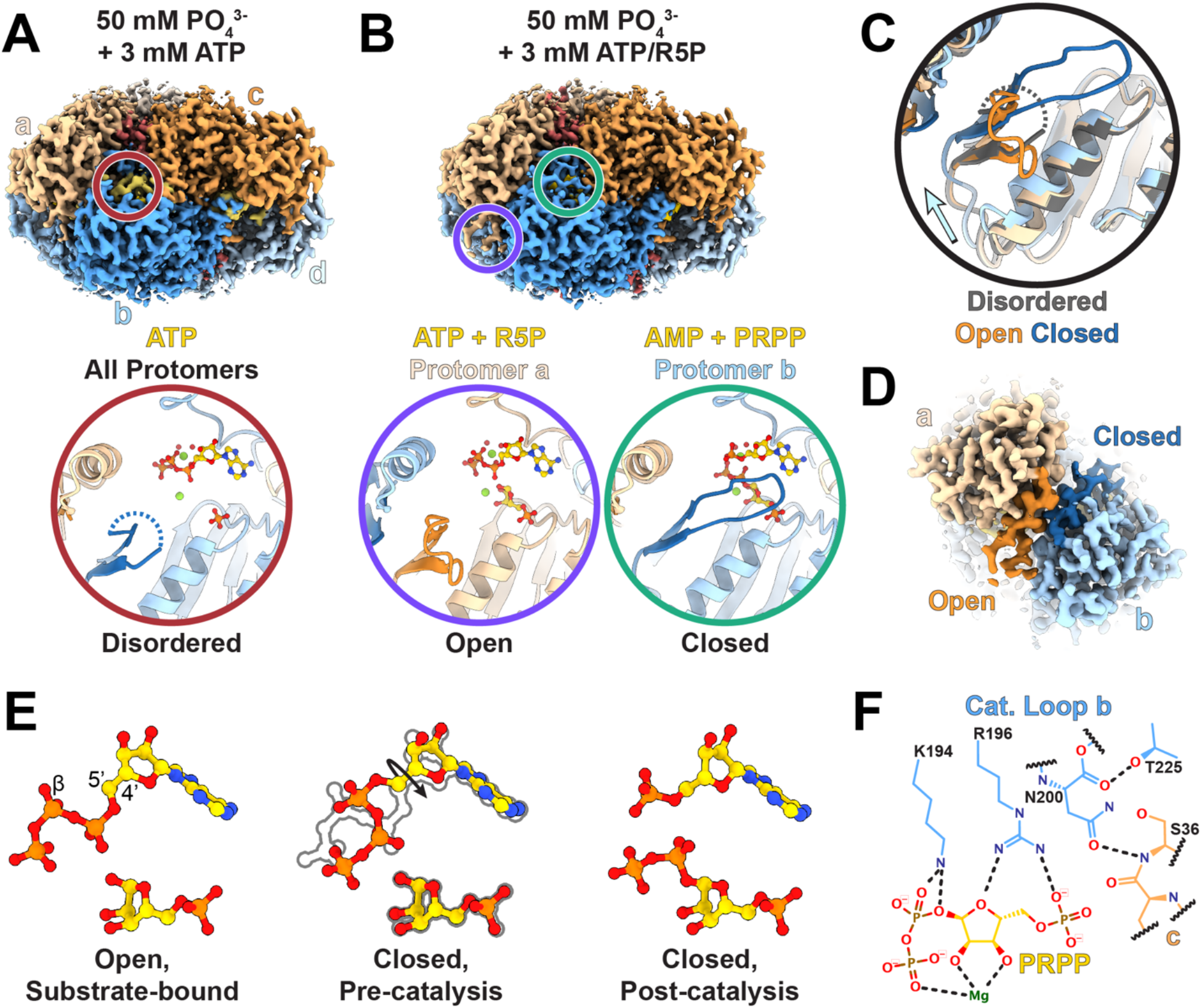
The allosteric interface coordinates catalysis across protomers. A. Volume of one hexamer from a filament of hPRPS1 bound to ATP and zoom in of active site indicated in by red circle including the catalytic loop (dark blue), ATP (yellow), phosphate, magnesium, and coordinated waters. B. Volume of one hexamer from a filament of hPRPS1 in the presence of ATP and ribose-5-phosphate and zoom in of neighboring active sites indicated by circles, with protomer a having the catalytic loop in an open position and bound to substrates (purple), and protomer b with the catalytic loop closed and bound to products (teal). C. Overlay of active sites shown in A and B, indicating shifts in the catalytic domain that accompany movement of the catalytic loop. D. Volume showing neighboring active sites with open (dark orange) and closed (dark blue) catalytic loops. E. Positioning of ligands in the substrate-bound, open active site and both pre-catalysis and post catalysis ligands that fit the volume in the closed active site. F. Interactions between the closed catalytic loop and the active site. Dotted lines indicate hydrogen bonds.

Despite nearly identical filament assembly interfaces, the filaments have distinct helical symmetries, with 88º rotation and 63 Å rise per hexamer for the phosphate-bound structure and 94º/61 Å for the ADP-bound (Figure 2B). The differences in helical symmetry arise from conformational differences within the hexamer, essentially rigid body movements of protomers relative to each other, with rearrangement of the allosteric loop (residues 97-113) around the ADP/phosphate site (Figure 2E). All published crystal structures of wild-type human PRPS contain sulfate in the allosteric site, and not surprisingly, these structures best align with the phosphate-bound PRPS hexamer (Ext. Data Fig. 3D), whereas the ADP-bound hexamer more closely resembles *E. coli* hexamers (PDB IDs 4S2U and 6ASV) which do not contain ligands in the allosteric site. Density for the ligands is well resolved in both structures (Figure 2F), and differences in their binding interactions generate shifts in the position of the protomers relative to each other. Individual protomers in the phosphate-versus ADP-bound structures are nearly identical (RMSD 0.6 Å), but the orientation of the protomers within the hexamer changes (Ext. Data Fig. 3E). These rotational shifts are modest across the bowed dimer interface (a-f) but are more pronounced across the allosteric site (a-b and a-c). The net effect is that bowed interface dimers rotate as a unit relative to neighboring dimers, resulting in a more open phosphate-bound filament and a more compact ADP bound filament (Supplementary Movie 1).

### Active PRPS1 structures reveals coordinated catalytic domain movements

We next asked how the presence of substrates affects the PRPS1 filament structure. We determined three structures, all in the presence of phosphate: one with ATP alone (2.2 Å resolution), one in the presence of ATP and ribose-5-phosphate while actively turning over these substrates (2.0 Å), and one with the product PRPP (2.4 Å) (Figure 3A-B, Ext. Data Fig. 4A-B, Table 1). In all three filaments phosphate is clearly visible bound in the allosteric site, and the conformation of the regulatory loop, the relationship of neighboring protomers, and the filament assembly interface are nearly identical to the phosphate-only structure. Thus, there are not large changes in the filament architecture upon substrate binding or during active catalysis.

**Figure 4.**
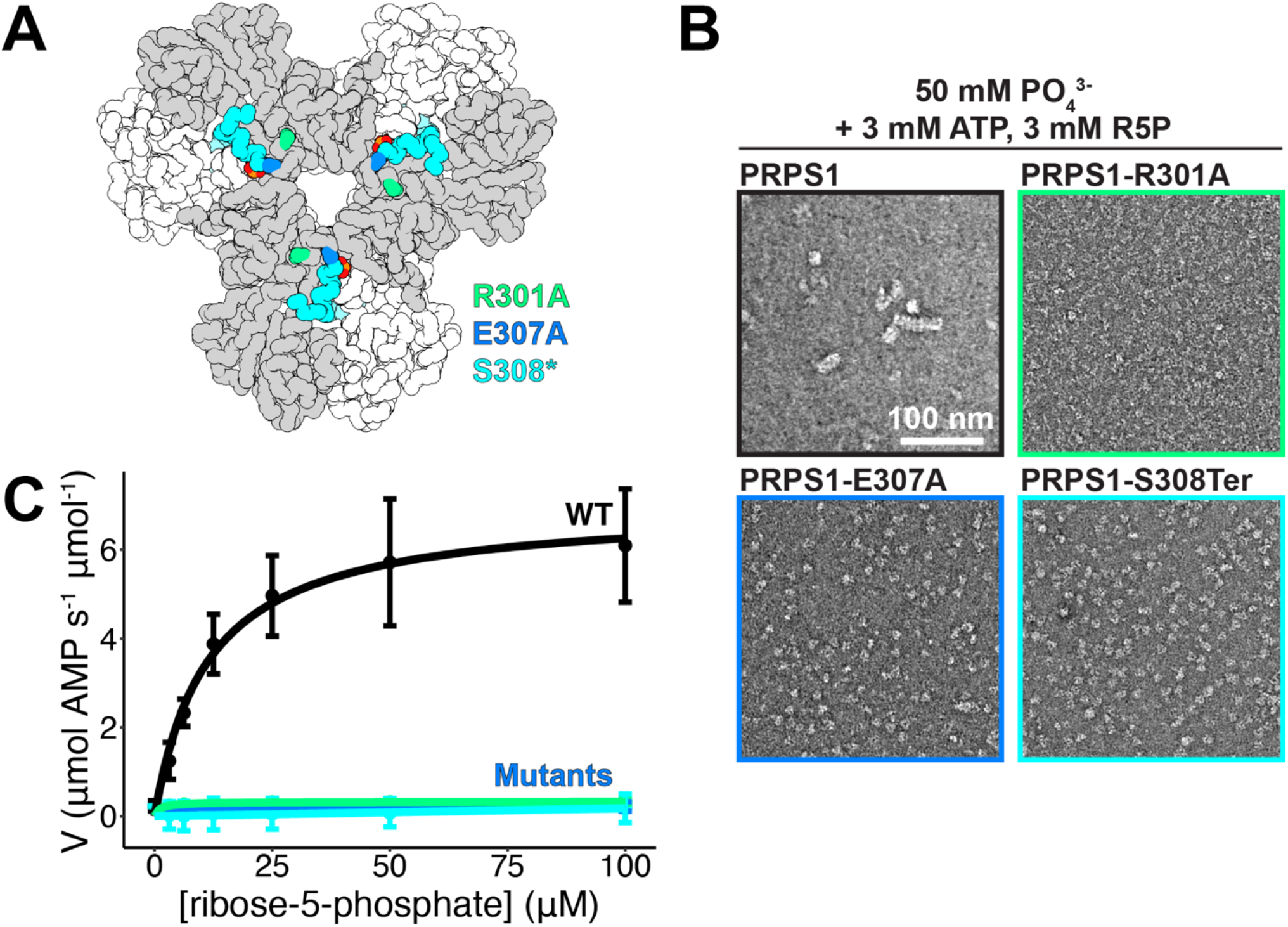
Mutation of filament interface residues decreases catalysis. A. Locations of the three engineered, filament-interface mutations: R301A (green), E307A (blue), and S308* (cyan). B. Negative stain EM images of the wild-type enzyme and the three mutations in the presence of phosphate, ATP, and ribose-5-phosphate. C. Substrate kinetics at equimolar protein concentration showing the catalytic activity of the wild-type protein and the three mutants.

One major structural difference, though, was clear in the ATP/R5P-bound structure, where additional disordered density near the active site suggested potential conformational variability during the catalytic cycle. The catalytic loop (residues 196-202) of the ATP-bound structure, like the phosphate-only structure, is not resolved in the map, but when ribose-5-phosphate is added there appeared to be additional poorly-ordered density around the active site, which is also seen in the structure containing only PRPP (Figure 3A, Ext. Data Fig. 4D-E). To better resolve the ATP/R5P active site structure, we used symmetry expansion and classification focused on a single protomer, followed by local refinement of the hexamer (Ext. Data Fig. 2 and 5A, Table 1). This resolved the catalytic loop in distinct opened and closed conformations (Figure 3B, Ext. Data Fig. 4C-E). In the open state, the overall protomer conformation is identical to the ATP or phosphate alone structures, the difference being that the catalytic loop is fully resolved in an extended conformation. In the closed state, the short pair of beta strands at the base of the catalytic loop rotates 11º, and the catalytic loop itself rearranges and flips about 45º to extend into the active site and contact the substrates. These changes near the active site are accompanied by a 5.7º rotation of the catalytic domain relative to the allosteric domain (Figure 3C, blue arrow, Ext. Data Fig. 4F).

**Figure 5.**
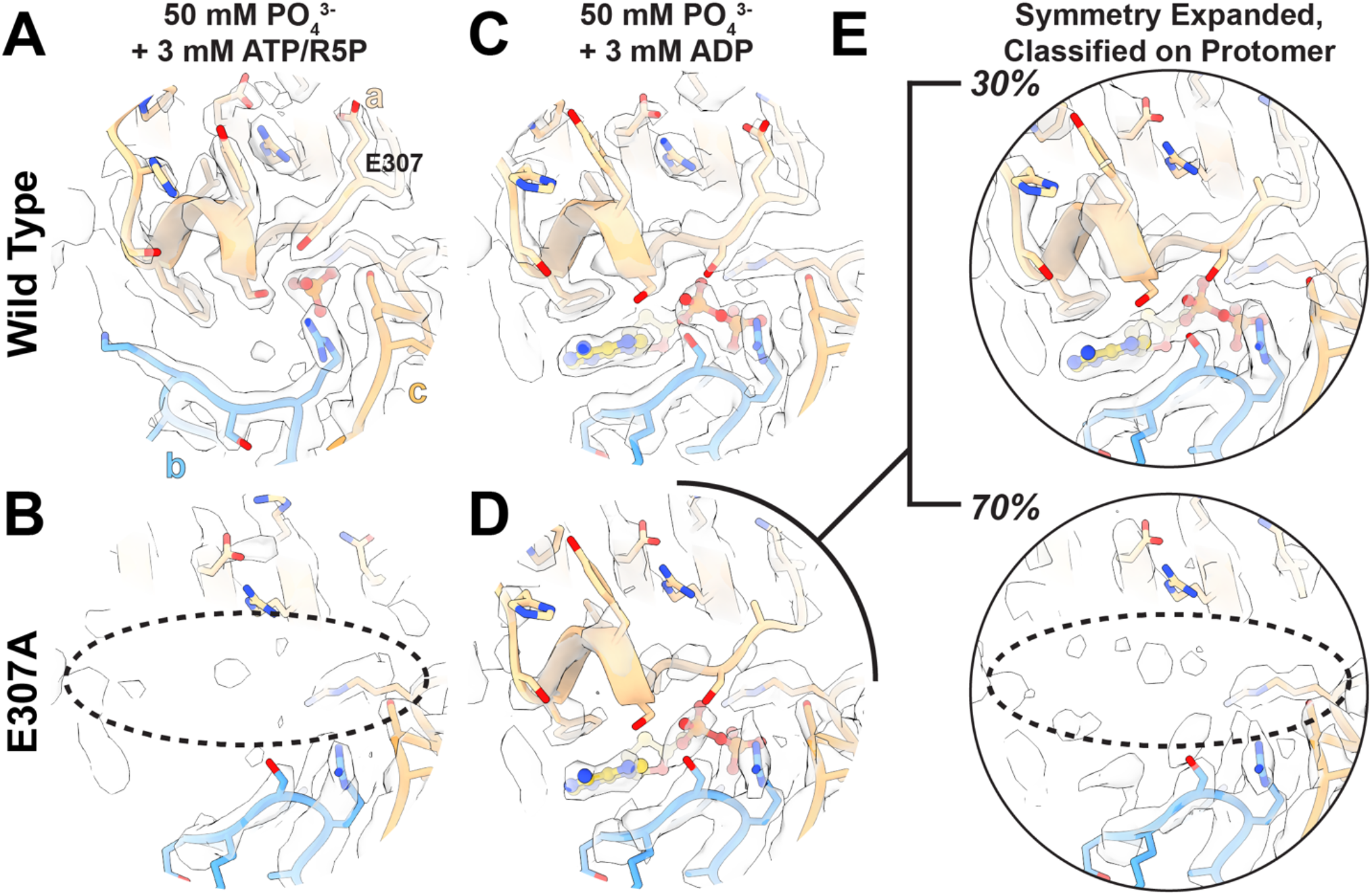
Filament formation stabilizes the C-terminus and allosteric site. A. The allosteric site and C-terminus of PRPS1 in the presence of phosphate, ATP, and ribose-5-phosphate. B. The allosteric site and C-terminus of PRPS1-E307A. C. The allosteric site and C-terminus of PRPS1 in the presence of phosphate and ADP. D. The allosteric site and C-terminus of PRPS1-E307A in the presence of phosphate and ADP. E. The PRPS1-E307A dataset was symmetry expanded and classified without alignment on a masked protomer, and two volumes were locally refined. *Top*. Map and model of the PRPS1-E307A showing the presence of the C-terminus and ADP bound in the allosteric site. *Bottom*. Map and model of the PRPS1-E307A lacking ADP in the allosteric site and a bound C-terminus.

Surprisingly, the conformations of neighboring protomers are linked. Our data processing strategy, with local refinement of orientations for entire hexamers after classification on a single protomer, revealed that when the catalytic loop of protomer a is closed, the catalytic loop of protomer b is open, and vice versa (Figure 3D, Ext. Data Fig. 4G). This relationship does not hold across the other, “bowed” interface with protomer c, as the catalytic loops in the active sites of protomers c through f are not well resolved. This suggests that the movements are anti-correlated within a-b dimers but uncorrelated between bowed dimers. Similarly, there does not appear to be coordination of active site states along the filament. The coordination between protomers is mediated by interactions between the short beta strands at the base of each catalytic loop, which remain in contact in the switch from open to closed. This suggests a possible “reload” mechanism for the active site, where closing of one catalytic loop pulls open the neighboring protomer, allowing catalysis in the first site and exchange of products for substrates in the neighbor (Supplementary Movie 2).

The density of the active site ligands differs in the open and closed conformations, suggesting rearrangements of the substrates accompany closure of the catalytic loop. When ATP alone is bound in the active site, the β-phosphate that is transferred to ribose-5-phosphate points out of the active site (Ext. Data Fig. 6A). In the open-loop conformation, the ATP and ribose-5-phosphate are bound in the active site, with ATP in the same pose as the ATP alone structure, suggesting a substrate bound pre-catalysis state (Figure 3E, Ext. Data Fig. 6B). When the catalytic loop is closed, the cryo-EM map supports placement of either substrates (ATP and ribose-5-phosphate), products (AMP and PRPP), or possibly a mixture of the two (Figure 3E, Ext. Data Fig. 6C-D). While the ribose and the adenosine moiety are in nearly identical poses in both open and closed states, the phosphates from ATP are reoriented. In the closed conformation the β-phosphate of ATP points into the active site, towards the 1’ carbon in ribose-5-phosphate, which requires an approximately 180° turn of the 4’-5’ bond in ATP relative to the pose in the open state and places it into a pre-catalysis position (Figure 3E). This position of pyrophosphate overlays with the position of the pyrophosphate group in PRPP in a separate structure with PRPP alone (Ext. Data Fig. 6E). The structures, then, support a model where positioning of the catalytic loop into the active site changes the position of the phosphates in ATP, a likely prerequisite for catalysis.

**Figure 6.**
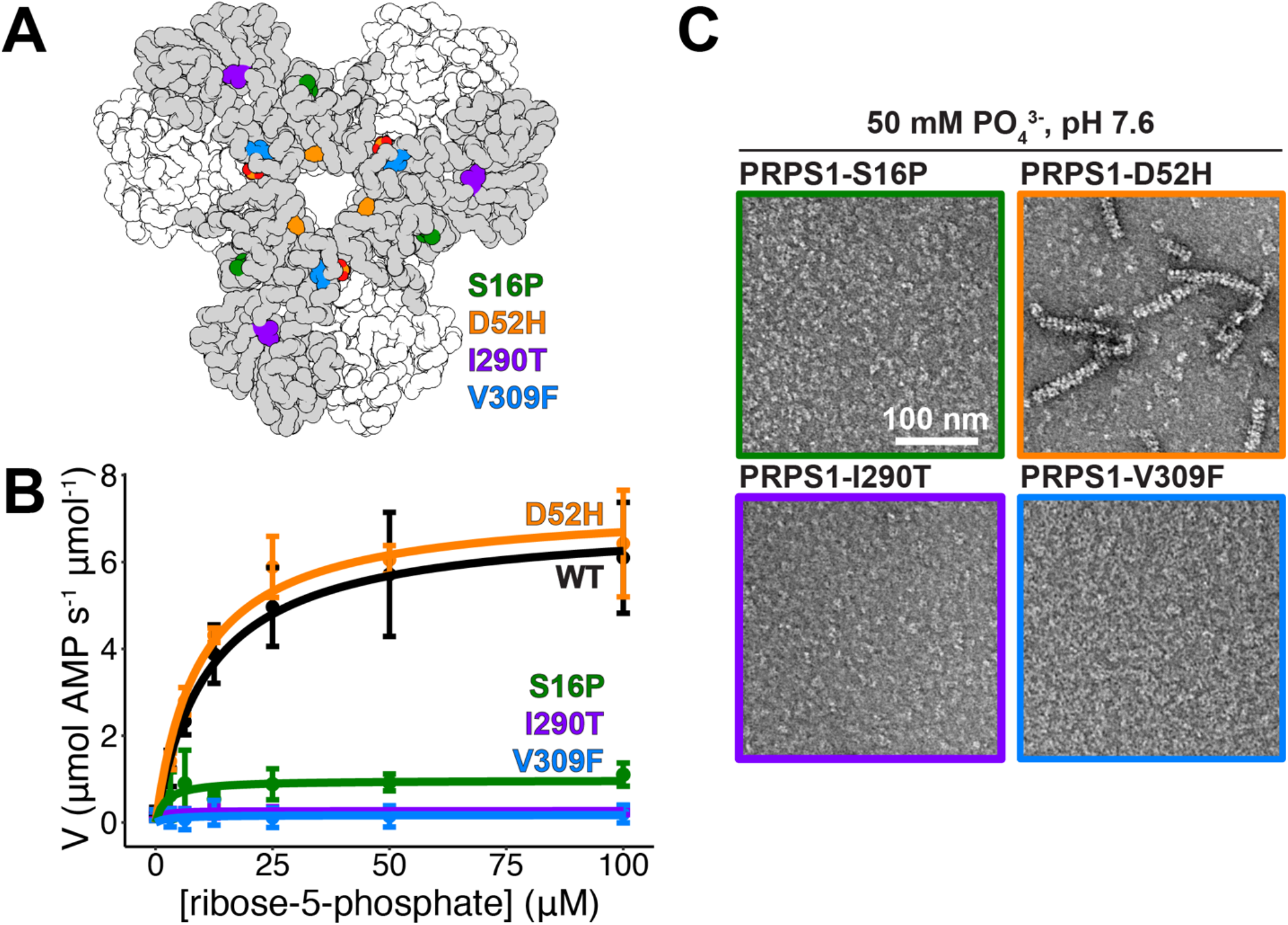
Mutations near the N- and C-termini alter filament formation which correlates with catalysis. A. Locations of the four mutations that cause disease: S16P, D52H, I290T, and V309F. B. Negative stain EM images of the wildtype enzyme and the three mutations in the presence of phosphate. C. Substrate kinetics of the wild type protein and the four mutations at equimolar protein concentration.

The position of residues in the catalytic loop provides a framework for understanding their functions in catalysis (Figure 3F). In the closed position the catalytic loop primarily contacts ribose-5-phosphate/PRPP, likely explaining why the loop remains disordered in the ATP-alone structure. In the closed state, R196 interacts with the 5’ phosphate and the ribose oxygen, suggesting it senses the presence of the substrate. N200 likely stabilizes the catalytic loop by interactions within the active site and with a neighboring protomer. These interactions position K194 to form hydrogen bonds with the ATP β-phosphate, suggesting it plays a direct role in catalysis; this interaction likely provides the basis for the catalytic loop-induced rearrangement of the ATP phosphates.

There are three existing PRPS crystal structures that contain a resolved catalytic loop in the closed position; two structures of PRPS from *Thermus thermophilus* (5T3O and 7PN0; hexamers) and one from *Thermoplasma volcanium* (3MBI; dimer)^18,31^. When aligned with the closed-loop PRPS1 protomer, the loops and the arginines and lysines occupy a similar position, despite the absence of ribose-5-phosphate in the active site (Ext. Data Fig. 4H). Moreover, the *T. thermophilus* hexamer structures have the same anti-correlated open-closed relationship between protomers that we observe in the human enzyme (Ext. Data Fig. 4I). Together, this suggests that the “reload” mechanism is a deeply evolutionarily conserved element of the catalytic cycle in hexameric PRPS.

### Disruption of the filament interface decreases catalysis in PRPS1

We next asked whether PRPS1 filament assembly has a functional role at the level of enzyme activity. We mutated residues in the filament interface to prevent filament assembly (Figure 4A). Mutation of either R301 or E307 to alanine, or truncation of the C-terminus at S308, prevents assembly under conditions where wild type PRPS1 forms filaments, including in the presence of phosphate, phosphate and substrates, and the inhibitor ADP (Figure 4B, Ext. Data Fig. 7A-B). Enzyme assays show that disruption of filament formation decreases catalysis (Figure 3C, Ext. Data Fig. 8), with turnover not detectable in conditions identical to the wild type protein. Substrate kinetics reveal a large decrease in K_cat_ for each of the mutants (Ext. Data Fig. 7C-D), indicating that substrate turnover is severely affected. This suggests that filament assembly plays a critical role in increasing the intrinsic catalytic activity of PRPS1.

**Figure 7:**
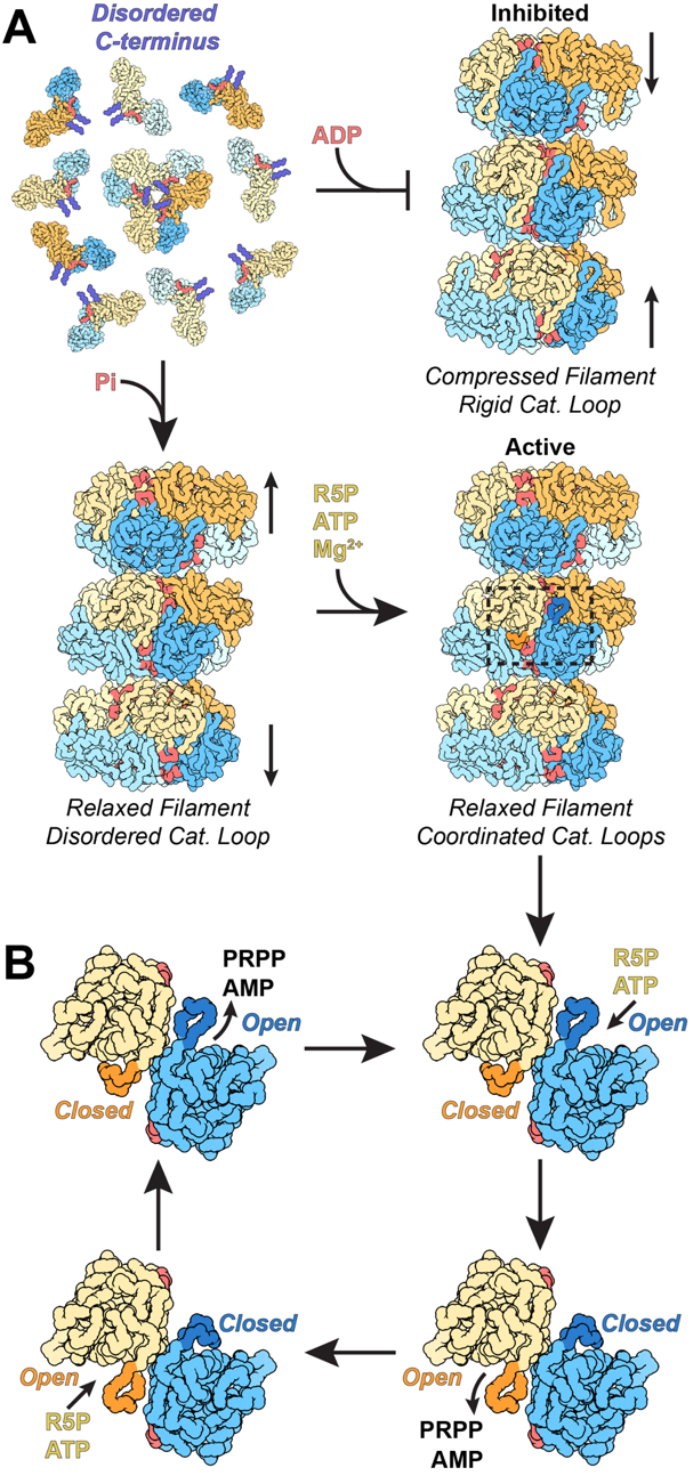
PRPS1 Filaments stabilize the allosteric site, reinforcing the inhibited and active conformations and facilitating a catalytic reload mechanism. A. In solution, PRPS1 assembles into dimers and occasional hexamers, where the C-terminus is disordered (purple). When ADP is added, the protein oligomerizes into compressed filaments, stabilizing the C-terminus and facilitating inhibition by locking the catalytic domains into a rigid conformation. When phosphate is added, the C-terminus is also stabilized, but the protein assembles into a more relaxed filament, where the catalytic domains of each protomer can flex. Addition of magnesium, ATP, and ribose-5-phosphate start a cycle where the paired catalytic domains are anti-correlated, promoting a reload mechanism. B. ATP and ribose-5-phosphate bind in an active site with an open catalytic loop (blue protomer). Binding of ribose-5-phosphate triggers loop closure (dark blue), which leads to the rearrangement of ATP within the active site, followed by catalysis. Closure of one catalytic loop triggers opening of the neighboring catalytic domain (orange protomer) and loop (dark orange), which releases AMP and PRPP. Binding of the second set of substrates triggers the same sequence of events, closing the loop (orange) and opening its neighbor (blue). This “reload” mechanism facilitates rapid catalysis.

### PRPS1-E307A filament interface mutant adopts an inhibited conformation in the hexamer

To characterize the mechanism by which filament assembly enhances activity, we determined structures of PRPS1-E307A in ADP- and substrate-bound free hexamers at 2.4 Å and 2.5 Å resolution, respectively (Ext. Data Fig. 2, 5, and 9, Table 1). In the substrate-bound PRPS1-E307A hexamer the allosteric site has no density for phosphate and the entire C-terminal region that contributes to the allosteric site (residues 306-315) is disordered (Fig. 5A, Ext. Data Fig. 9A).

Moreover, in the absence of bound phosphate the overall hexamer is in the inhibited conformation (Figure 5B), despite clear density for substrates in the active site (Ext. Data Fig. 5B). Additionally, there is no evidence for closure of the catalytic loop. These dramatic differences from the substrate-bound wild type enzyme, despite identical ligand conditions, likely explain why the mutant has reduced activity.

In contrast to the active state, PRPS1-E307A + ADP is broadly similar to the wild type enzyme, with density for the allosteric site and residues 306-315. However, the map suggested that there was only partial occupancy of the allosteric site (Ext. Data Fig. 9B). After symmetry expansion, classification by protomer, and local refinement, the map resolved into two classes, one with the allosteric pocket formed and occupied by ADP (Figure 5C, Ext. Data Fig. 9C) containing 30% of the particles in the dataset, and a second empty pocket with residues 306-315 disordered (Figure 5D; Ext. Data Fig. 9D) containing about 70% of the particles in the dataset.

The structures of filament-incompetent PRPS1-E307A in two different ligand states demonstrate that outside of the filament the PRPS1 allosteric site is destabilized. This suggests that C-terminal residues 306-315 are relatively weakly bound to the core of the enzyme and become stabilized by interaction with allosteric ligands. The participation of E307 in the filament interface likely anchors the C-terminal region, which in the context of the filament is also sterically constrained from sampling the disordered state observed in the free PRPS1-E307A hexamers. Thus, filament assembly contacts act to stabilize the allosteric site, enabling binding of the essential activator phosphate, which shifts hexamer conformation to promote activity.

### Disease-associated PRPS1 mutations are defective in filament assembly and catalysis

Many point mutations in PRPS1 that are associated with disease are far from the active site but lead to changes in catalytic activity. Both gain-of-function and loss-of-function mutations have been identified, associated with a spectrum of diseases caused by over- or underproduction of nucleotides. As disruption of filament formation has severe consequences on catalytic activity, we explored the effects of a subset of disease mutations that are located near the filament assembly interface (S16P, D52H, I290T, V309F) (Figure 6A)^32–35^.

Substrate kinetics of purified protein followed the same pattern of activity as described in the literature, where whole cell lysate has been used to assess activity^33–37^. D52H, a gain-of-function mutant that is more active in low phosphate conditions, has a V_max_ similar to the wild type enzyme in the conditions tested, where phosphate is not limiting (Figure 6C, Ext. Data Fig 10A)^32^,. The other three mutations are all loss-of-function mutations, and all three have decreased or unmeasurable V_max_ values (Figure 6B, Ext. Data Fig. 10A).

The pattern of activity displayed by the mutations is paralleled in their assembly into filaments. The D52H mutation retains filament assembly similar to the wild type enzyme (Figure 6C, Ext. Data Fig. 10 B-C). Conversely, all of the loss of function mutations are defective in filament assembly. Like the engineered, filament-disrupting mutations described above, this suggests that catalytic activity is tied to filament formation, and disruption of the filament interface decreases the activity of the enzyme.

## Discussion

Regulation of PRPS has long been interpreted in the context of in its hexameric form — despite early observations that hexamers reversibly assemble into higher order oligomers, or “aggregates,” the function of these larger structures has largely gone unexplored^8,12–14^. Recent observations that PRPS forms micron-scale filaments in cells of diverse species has renewed interest in the functional properties of the higher order oligomers^19–21^. Here, we have shown that the human enzyme PRPS1 forms filaments in the presence of the allosteric effectors phosphate and ADP, and that the filaments are much more active than free hexamers. Increased activity in the filament likely arises from stabilization of the allosteric regulatory site by filament assembly contacts; in free hexamers the allosteric site is disordered, preventing binding of the essential activator phosphate and the consequent conformational changes that increase catalytic activity. Assembly into filaments, therefore, provides an additional layer of regulatory control on top of established allosteric mechanisms within the hexamer, to tune PRPS1 enzymatic activity (Figure 7A). Our in vitro characterization of the filament assembly-based regulation mechanisms lays the groundwork for future studies to probe the function of the polymers in the cellular context.

The link between filament assembly and increased activity also holds with point mutants near the filament assembly interface that are associated with human diseases. Mutations in PRPS1 fall along a spectrum from overactive to underactive^4^. Most of these mutations are found outside of the active site and do not produce as severe catalytic defects as the engineered filament mutations. However, as filament formation is crucial for efficient catalysis, even a slight shift in the propensity to assemble could have serious consequences for catalytic activity. This becomes even more pronounced in the male population, as PRPS is located on the X-chromosome^38^. It appears that in some cases, disease mechanisms may be linked to disruption of filament assembly, suggesting an important role for PRPS1 filaments in human health.

The deep evolutionary conservation of the residues that mediate PRPS1 filament assembly suggests that PRPS likely assembles filaments in most species. Indeed, fluorescence imaging has shown that PRPS assembles into micron scale filamentous structures in human, rat, drosophila, budding yeast, and *E. coli* cells^19–21^. Moreover, a recent study showed that PRPS from *E. coli* assembles filaments in vitro in both the active and the inactive state^19^. In this case, it appears that the *E. coli* protein forms two distinct filament interfaces depending on the ligand bound in the allosteric site. The filament assembly interface in the inhibited *E. coli* filament appears to be nearly identical to the assembly contacts seen in all the activity states of human PRPS1 filaments, supporting the idea that filament formation has been conserved through billions of years of evolution, although the functional consequences of assembly may have evolved.

Visualizing active PRPS1 at high resolution while in the process of turning over substrates provides insights into its catalytic mechanism (Figure 7B). We found that opening and closing of the catalytic loop is anticorrelated in neighboring protomers of the a-b dimer, so that closure at one site is coupled to opening of the adjacent site, suggesting a reload mechanism that allows alternating catalysis and release. The closed state is largely stabilized by interactions of Arg196 with ribose-5-phosphate so that closure is unlikely in the absence of substrate, which would prevent unproductive turnover of ATP. Closure of the catalytic loop induces rearrangement of the ATP phosphates to position the β- and γ-phosphates for transfer to ribose-5-phosphate, and positions the residue Lys194 opposite the β-phosphate to support catalysis. The catalytic loop must open post-catalysis to release the PRPP product, with opening likely driven by conformational changes in the adjacent site upon substrate binding, generating a back-and-forth cycle of opening and closing in adjacent active sites. During this cycle the allosteric domain core of each hexamer remains fixed, and the movements are only of the adjacent catalytic domains relative to this core, explaining the coordination of binding at a-b dimers but no apparent coordination with other protomers in the hexamer. The reload mechanism also appears to be broadly conserved throughout domains of life, as the open/closed pairing appears in several crystal structures.

Taken together, the very low activity of engineered or disease mutations that disrupt assembly would suggest that PRPS1 must be assembled into filaments for robust activity, although transient association of hexamers might allow for low levels of activity. While little is known about the conditions that promote PRPS1 assembly in cells, controlling cellular assembly may provide a powerful mechanism for regulating activity. Similar mechanisms have been shown for other metabolic enzymes. In one striking example, IMP dehydrogenase, which has increased activity in filaments assembles micron-scale filaments in activated T-cells in response to metabolite levels and signaling through calcium levels and MTOR activity^29,39,40^. The structural and in vitro biochemical characterization of PRPS1 filaments presented here provides a framework for understanding PRPS1 regulation in the more complicated cellular environment.

## Acknowledgements

We thank the Arnold and Mabel Beckman Cryo-EM Center at the University of Washington for electron microscope use. We also thank members of the Kollman group for valuable feedback provided during cryo-EM data collection and processing. This work was supported by the US National Institutes of Health (grant nos. R01GM118396 and S10OD023476 to J.M.K., 1F32AI145111 to K.L.H.)

## Methods

### Protein Expression and Purification

BL21 (DE3) pLysS *E. coli* were transformed with a 6xHisSUMO- PRPS1 wild type or mutant construct on a pSMT3-Kan vector, and were grown at 37°C in Luria broth until an OD_600_ of 0.6-0.8 was reached. Cultures were shifted to 16°C, induced with 1 mM IPTG, and grown overnight. Cells were pelleted stored at -20°C. For purification, pellets were resuspended in lysis buffer (50 mM HEPES pH 8, 400 mM NaCl, 75 mM Imidazole pH 8, 5% v/v glycerol, 5 mM 2-Mercaptoethanol, supplemented with Roche Protease Inhibitor Tablets) and lysed using an Emulsiflex-05 homogenizer. Lysate was cleared by centrifugation and the protein was purified by nickel affinity chromatography (Thermo Scientific HisPur Ni-NTA Resin). After applying cleared lysate to the resin on-column, the resin was washed (50 mM HEPES pH 8, 400 mM NaCl, 75 mM Imidazole pH 8, 5% v/v glycerol, 1 mM DTT) and eluted with wash buffer supplemented with an imidazole step gradient (imidazole concentrations: 0.2 M, 0.3 M, 0.5 M, 0.7 M). Eluted fractions were then incubated with Ulp1 (1:100 Ulp1:PRPS), concentrated (Millipore Amicon Ultra 10K MWCO), and further purified by size exclusion chromatography using a Äkta Pure system (Cytiva Life Sciences) and a pre-equilibrated, GE Superdex 200 gel filtration column (50 mM HEPES pH 8, 400 mM NaCl, 5% v/v glycerol, 1 mM DTT). Peak fractions were concentrated (Amicon Ultra 10K MWCO), snap frozen with liquid nitrogen, and stored at -80.

### Analytical Size Exclusion Chromatography

Purified wild type or mutant protein from a frozen stock was diluted to 30 uM in buffer (50 mM Potassium Phosphate pH 7.6, 6 mM MgCl_2_, 0.1 mM EDTA, 1 mM DTT). Proteins or standards (150 uL) were loaded onto a GE Superose 6 Increase 10/300 GL column using and AKTA Pure and run using manufacturer’s instructions. Curves were exported from Unicorn (Cytiva Life Sciences) and plotted in RStudio (v1.4.1103).

### Negative Stain Electron Microscopy

Protein samples were assembled at 10 uM with buffer and ligands at concentrations indicated in the results. The samples were then incubated at 37 degrees C for 2 minutes, diluted to 0.5 uM in matched solution, and applied to glow-discharged continuous carbon film grids. Grids were then washed in ddH_2_O three times and negatively stained using 2% uranyl formate. Samples were imaged using a FEI Morgagni operating at 100 kV and a Gatan Orius CCD camera with the software package Digital Micrograph (Gatan) or using a FEI Tecnai G2 Spirit operating at 120 kV and a Gatan Ultrascan 4000 CCD camera with the software package Leginon^41^.

### Cryo-Electron Microscopy

Protein samples were assembled at 7-30 uM with buffer and ligands at concentrations indicated in Table 1. Protein solutions were applied to glow-discharged, C-flat 2/2 or 1.2/1.3 holey-carbon EM grids (Protochips Inc.), blotted, and plunge-frozen in liquid ethane using a manual plunging apparatus at room temperature. Data collection was performed using an FEI Titan Krios transmission electron microscope operating at 300 kV (equipped with a Gatan image filter (GIF) and post-GIF Gatan K2 or K3 Summit direct electron detector) and an FEI Glacios (equipped with a Gatan K2 Summit direct electron detector) both using the software package Leginon^41^. Where indicated in the Table, the stage was tilted to 45 degrees for tilted data collection.

### Data Processing

Movies were aligned, corrected for beam-induced motion, dose-weighted, and binned (2x) using the Relion 3.1 implementation of MotionCor2; CTF parameters were estimated using CTFFind4^42–44^. Motion corrected micrographs were imported into cryoSPARC (v3) and particles were picked using Blob Picker^45^. For a subset of datasets (PRPS1 + ADP), particles were picked from an initial subset of images, 2D classified and then the full set of particles were picked using the 2D averages in the Template Picker. Two to three rounds of 2D classification were used to remove noise and poorly aligning particles, and the remaining particles were used in *ab initio* reconstruction and 3D refinement with D3 symmetry imposed. The particles selected and the model generated in 3D refinement in cryoSPARC were then exported to Relion, where additional rounds of 3D refinement were completed, interspersed with CTF refinement, masked refinement, Bayesian polishing, signal subtraction, and map sharpening with resolution estimation (FSC cutoff of 0.143). For a subset of datasets, the particles were symmetry expanded after Bayesian polishing and sent through a round of 3D classification without alignment followed by a round of local refinement and map sharpening with resolution estimation^46^. The final refined, unsharpened maps were exported to Phenix (v1.18) where density modification and additional resolution estimation was performed (FSC cutoff of 0.5)^47^. Directional FSCs were calculated using the remote processing server for 3DFSC^48^.

### Model Building

PDB ID 2H06 and 3S5J were used as initial models for the PRPS1 + ADP and PRPS1 + ATP datasets, respectively. Models were iteratively refined using a combination of ISOLDE (v1.2) in ChimeraX (v1.3), Coot (v0.9.4.1), and phenix.real_space_refine and phenix.validation_cryoem in Phenix^49–52^. The phenix.validation_cryoem tool was used to generate model statistics. Models from the ADP and ATP datasets were used as initial models for the additional datasets.

### AMP-Glo Activity Assay

Wild type or mutant protein from a frozen stock was diluted to 2x assay concentration (0.01 – 20 uM) into enzyme assay buffer (50 mM Potassium Phosphate pH 7.6, 6 mM MgCl_2_, 0.1 mM EDTA, 1 mM DTT, 0.1 mg/mL bovine serum albumin). ATP and ribose-5-phosphate were diluted to 2x assay concentration into enzyme assay buffer. Protein and substrates were mixed one to one and allowed to react for 1-5 minutes. Reactions were diluted if needed to bring the AMP concentration within the linear window of the AMP Glo assay (Promega). The AMP Glo assay was then performed according to the manufacturer’s instructions. Briefly, 5 uL of AMP Glo Solution I were added to 5 uL of the enzyme reaction and incubated for 1 hour. 10 uL of AMP Detection Solution (a 1:100 dilution of AMP Glo Solution II into Kinase Glo I mixed directly before addition) were added to the reaction, placed in the dark, and incubated for 1 hour. Luminescence values were read in triplicate on a Thermo Scientific Varioscan LUX plate reader. Unless otherwise noted, experiments were repeated for a minimum of 3 replicates. Luminescence values were averaged and background subtracted, and AMP concentrations were calculated by comparison to a standard of known AMP concentrations. Kinetic parameters were calculated in RStudio (v1.4.1103), using the packages “dplyr” (v1.0.9) and “drc” (v3.0-1)

### Figure Assembly

Electron microscopy images were prepared in Fiji (v2.1.0). Map and model figures were prepared in Chimera (v1.15) and ChimeraX (v1.3). Plots and graphs were prepared in RStudio (v1.4.1103) using the package “ggplot2” (v3.3.5). All figures were assembled in Adobe Illustrator CC (v26.0.1).

## Extended Data Figures

**Extended Data Figure 1.**
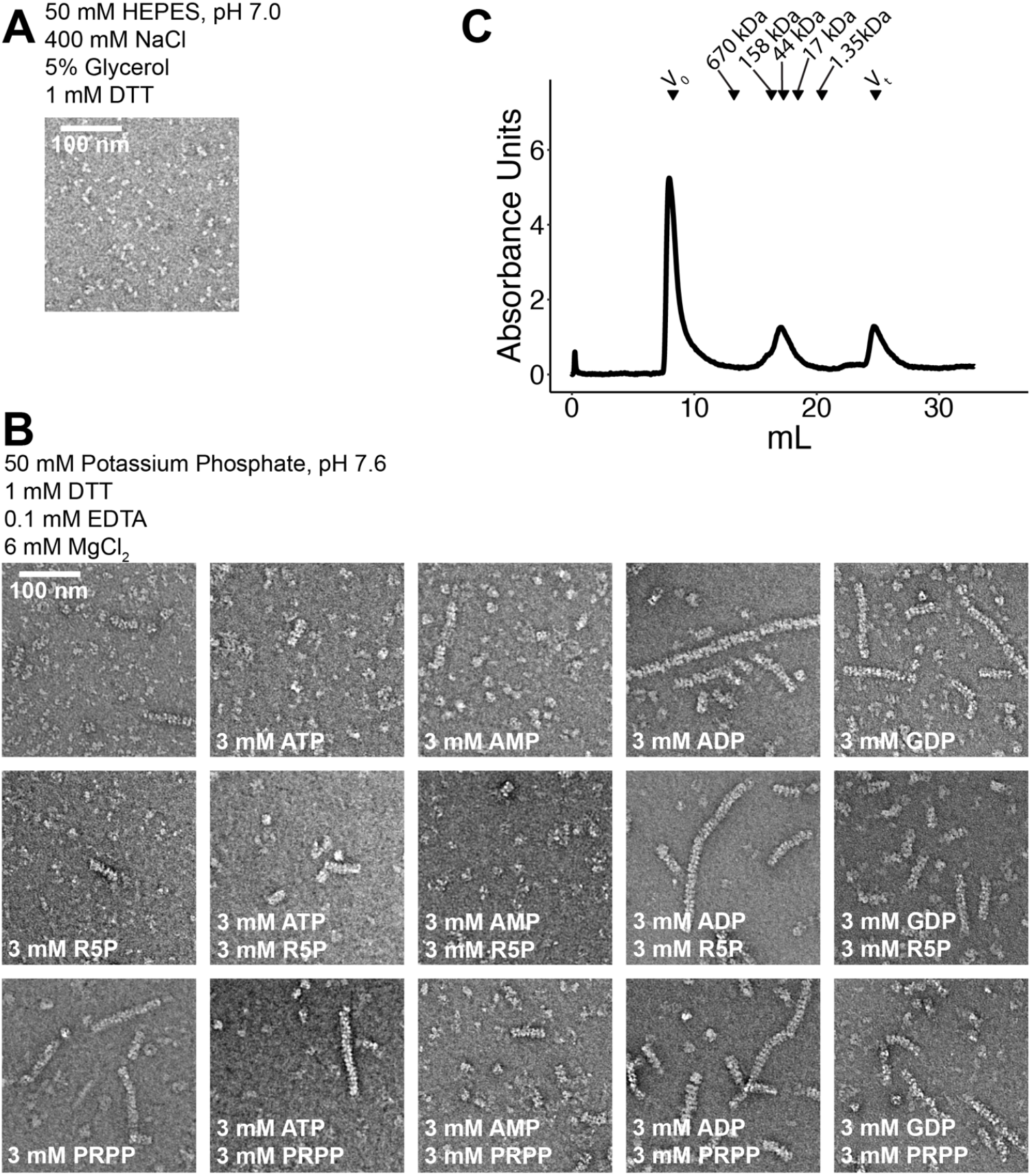
(Figure 2) Filament formation in PRPS1. A. Negative stain EM of purified PRPS protein in a HEPES/Salt buffer used for purification. B. Panel of negative stain EMs of PRPS1 in phosphate buffer in the presence of the indicated ligands. C. Elution profile from a size exclusion column (Superose 6) of PRPS1 in 50 mM phosphate buffer, pH 7.6.

**Extended Data Figure 2.**
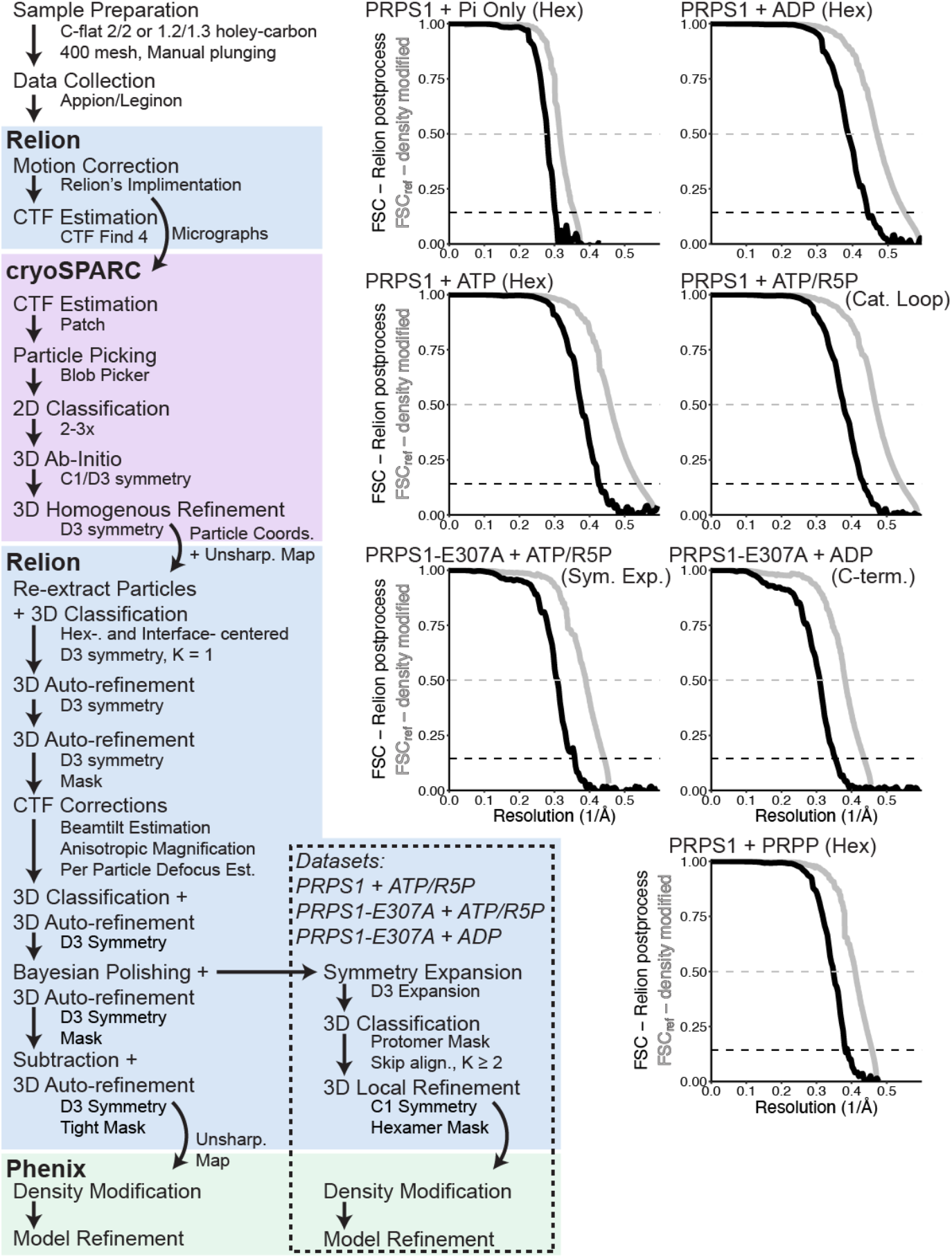
(Figure 2,3,5) Data processing and statistics for cryo-EM datasets. Left. Overview of the processing scheme for the datasets presented in this manuscript. Right. Fourier shell correlation curves calculated in Relion (black) and in Phenix (grey). One set of curves per dataset is shown. For full dataset statistics and information, see Data Table 1.

**Extended Data Figure 3:**
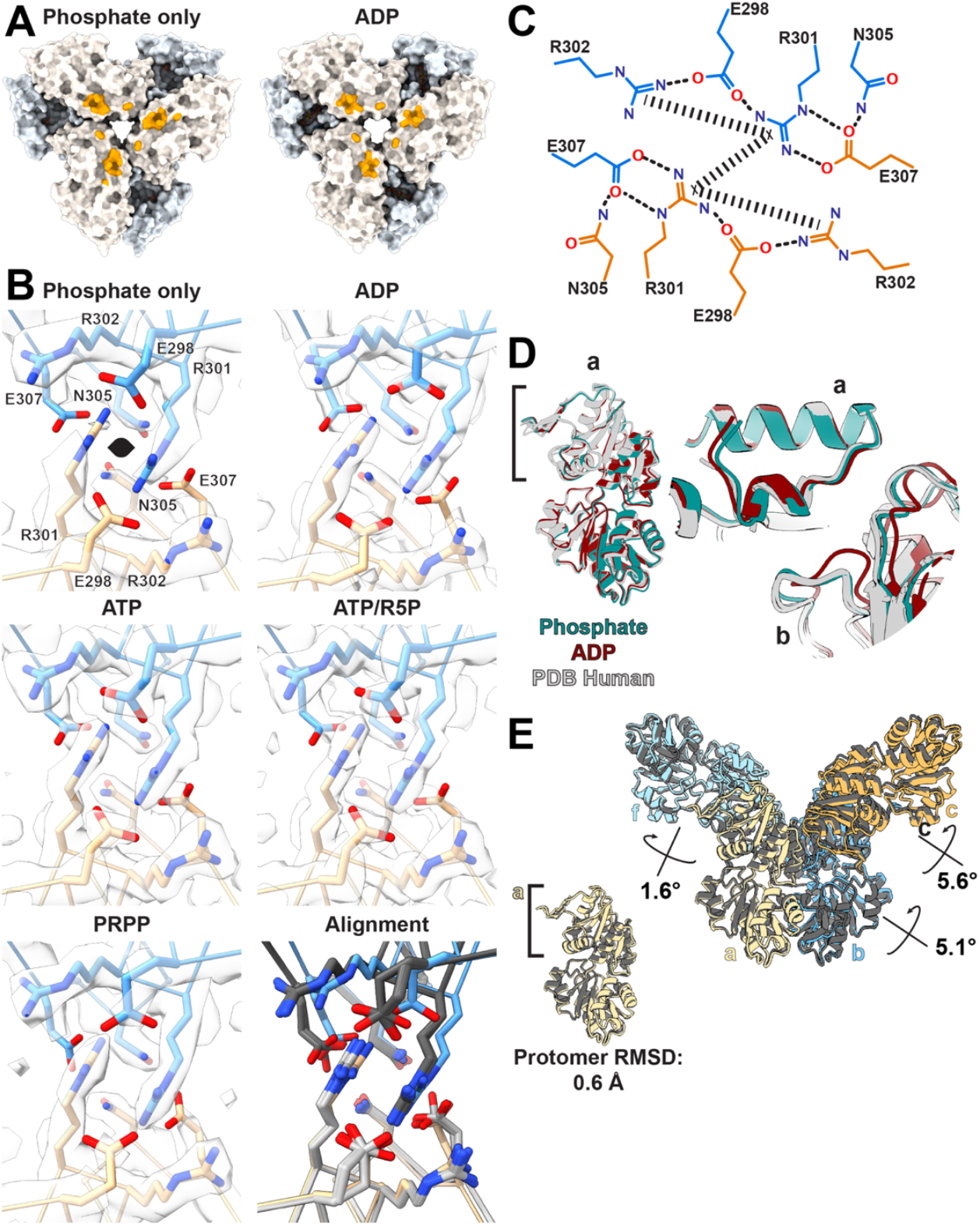
(Figure 2,3,5) Volumes and models of filament interface residues. A. Surface representation of filament interface in phosphate- or ADP-bound structures; orange patches indicate residues involved in the interface. B. Model and map of the primary interface residues in all filament structures presented in this manuscript. Bottom right panel shows the overly of the interfaces when aligned by the bottom protomer. ADP-bound structure colored in orange and blue; all others in grey. C. Schematic of primary interactions across the filament interface; rectangular dashed lines indicate pi-stacking interactions and rounded dashed lines indicate hydrogen bonds. D. Comparison of phosphate- (teal) and ADP-bound (maroon) structures to human crystal structures of wild type PRPS1 (grey). Structures have been aligned on the allosteric domain of protomer a. E. Alignment of the phosphate and ADP-bound structures on the allosteric domain (left) show minimal differences at the protomer level. Differences in the filaments arise from the orientations of the protomers relative to each other in the hexamers, with rotations of neighboring protomers relative to a as indicated.

**Extended Data Figure 4.**
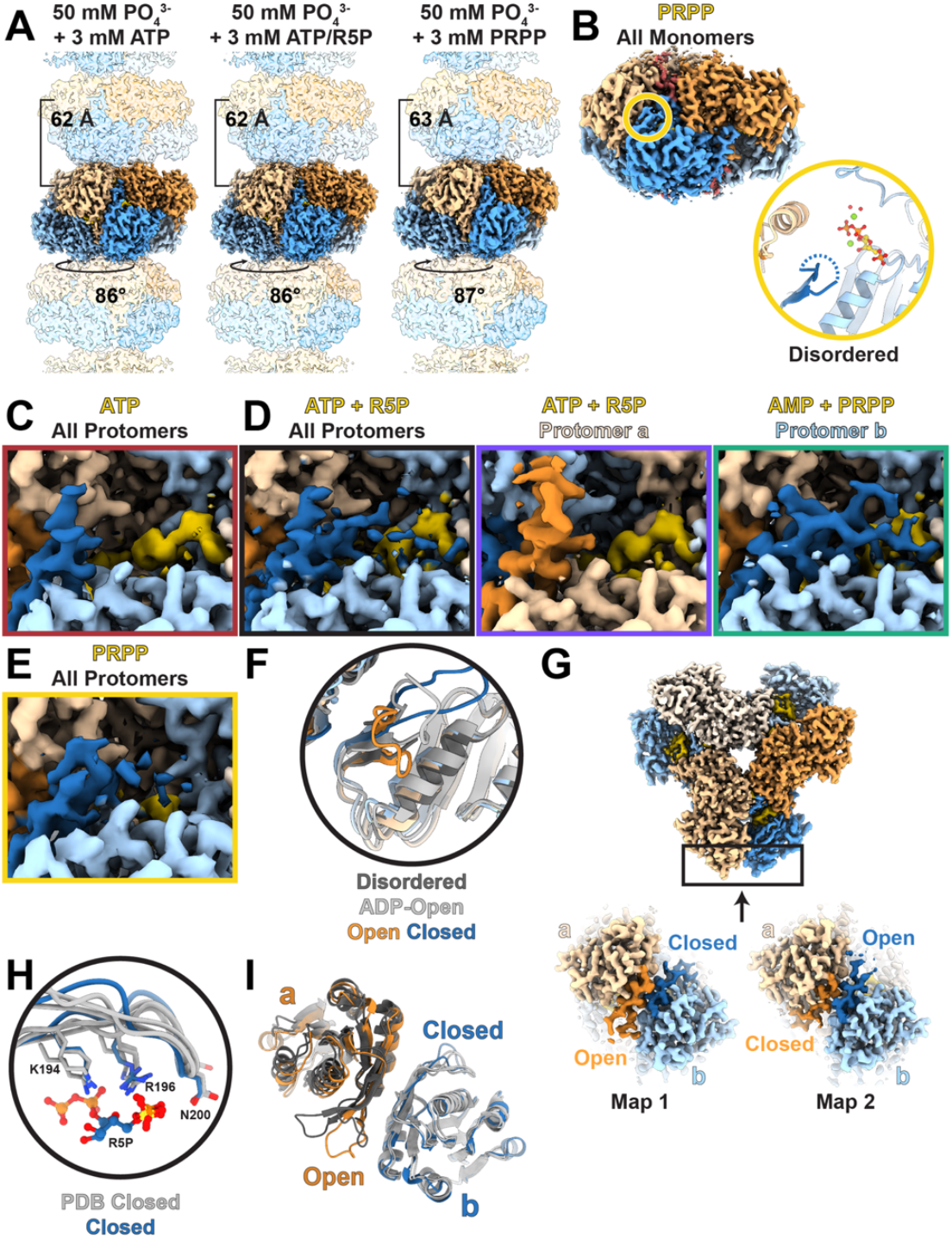
(Figure 3) Substrate- and product-bound filaments. A. Volume of PRPS1 filaments bound to phosphate/ATP (left), phosphate/ATP/R5P (middle), or phosphate/PRPP (right); protomers colored in blue or orange. B, top. Volume of one hexamer from a filament of PRPS1 bound to PRPP. Protomers are orange and blue, with the active site in yellow. B, bottom. Zoom in of active site indicated in (top), including the catalytic loop (dark blue), ATP (yellow), phosphate, magnesium, and coordinated waters. C-E. Volume showing the catalytic loop (dark blue or dark orange) and the ligands in the active site (yellow) for each of the filament structures presented in this work. F. Overlay of active sites shown in Main Text Figure B,C-D and also including PRPS1 + ADP (light grey). G. Volume (top) describing location of slices (bottom) showing catalytic domains in two maps with well-resolved catalytic loops. H. Overlay of PRPS1 + ATP/R5P closed catalytic loop and key residues from the three PRPS structures from the PDB that also contain a closed catalytic loop (3MBI from *Thermoplasma volcanium*; 5T3O and 7PN0 from *Thermus thermophilus*). PRPS1 with ATP/R5P in blue, PDB models in grey. I. Overlay of PRPS1 with ATP/R5P (blue/orange) and 5T3O and 7PN0 from *Thermus thermophilus* (greys), showing the neighboring open and closed catalytic loops.

**Extended Data Figure 5:**
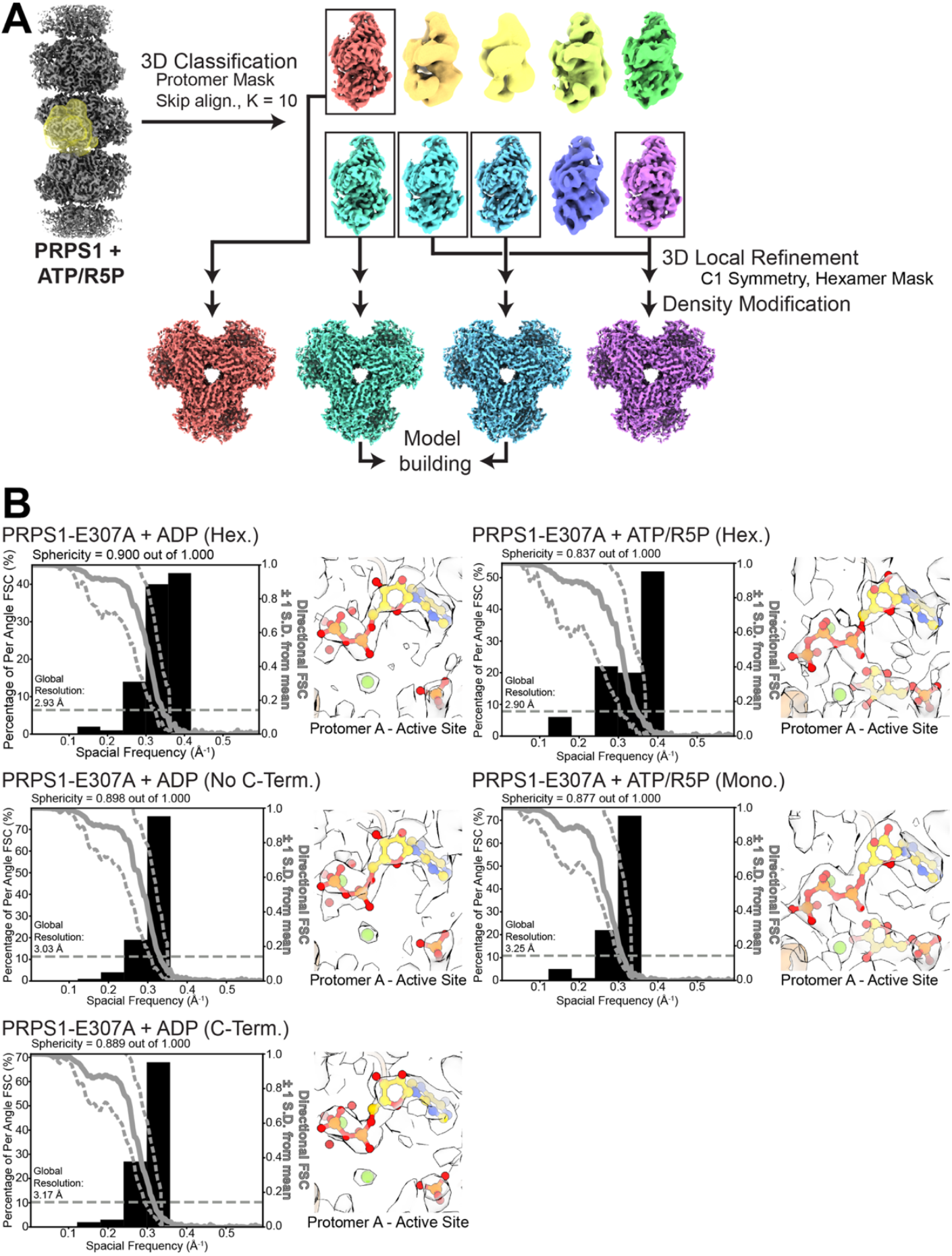
(Figure 3,5) Example classification scheme and directional FSCs. A. Classification scheme for PRPS1 + ATP/R5P after symmetry expansion. Particles were classified into ten classes, without alignment using a protomer mask. A subset of the resulting volumes was locally refined using a hexamer mask and exported to phenix for density modification. Two volumes were then used for model building. B. Directional FSC for volumes derived from tilted datasets and volumes and models from the active sites from protomer a of each map.

**Extended Data Figure 6.**
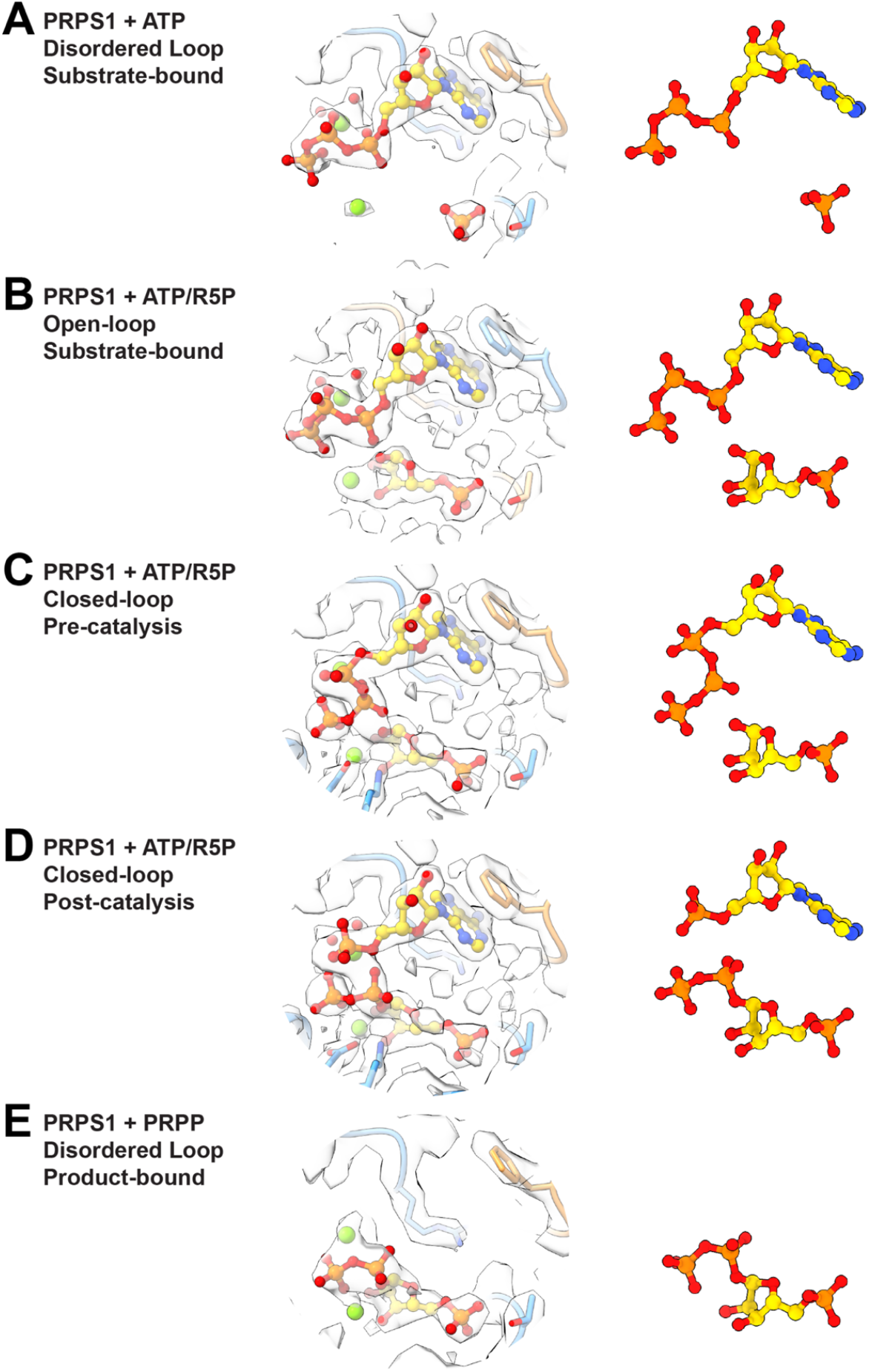
(Figure 3) Ligand volumes from substrate- and product-bound filaments. Panels show the volume and the ligands for (A) PRPS1 + ATP, (B) PRPS1 + ATP/R5P with open loop, (C) PRPS1 + ATP/R5P with closed loop, (D) PRPS1 + AMP/PRPP with closed loop, (E) PRPS1 + PRPP.

**Extended Data Figure 7.**
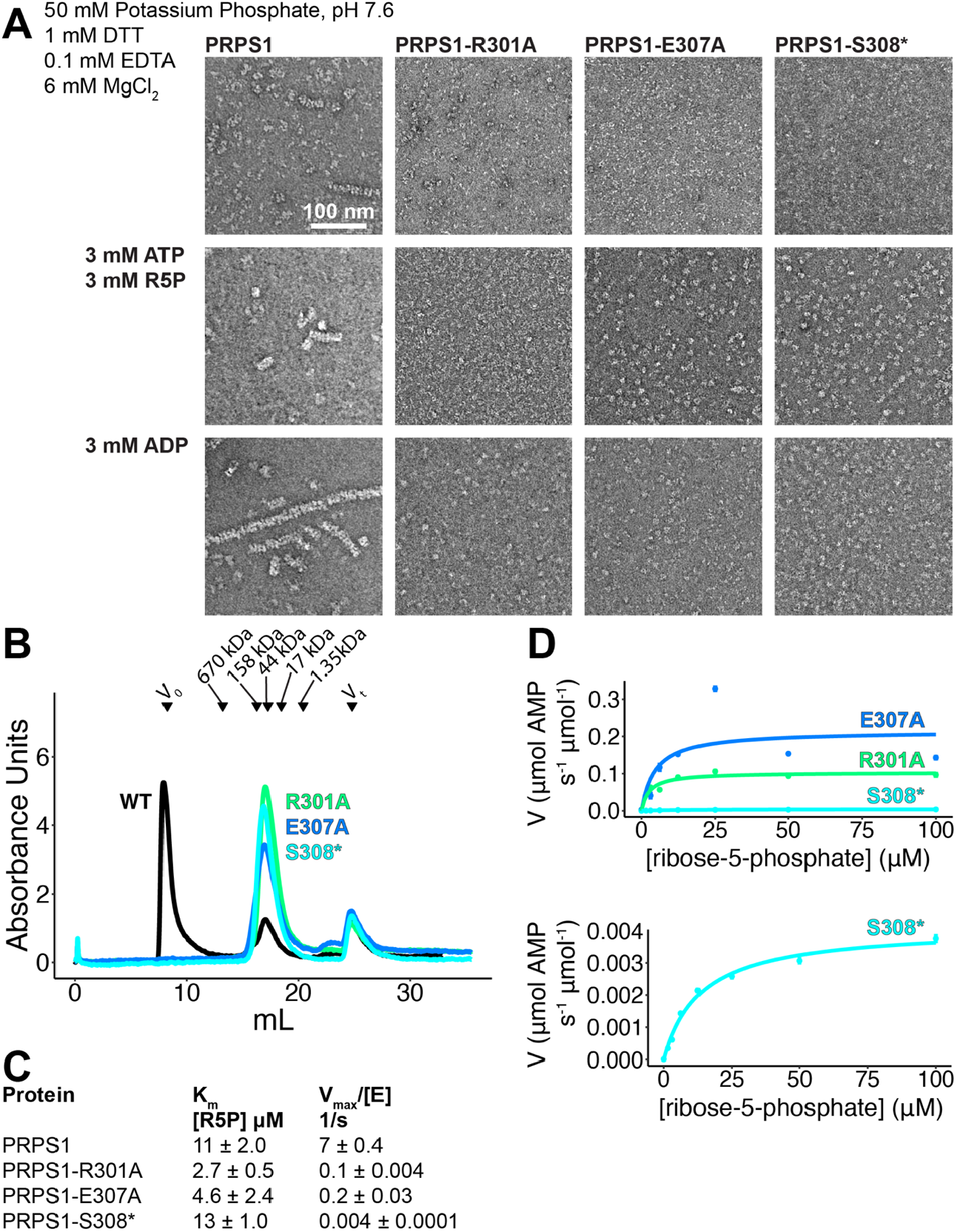
(Figure 4) Mutation of filament interface residues. A. Panel of negative stain EM of PRPS1 engineered mutations in phosphate buffer in the presence of the indicated ligands. B. Chromatography curves from a Superose 6 of PRPS1 and three engineered, filament-interface mutations. C. Kinetic parameters for the wild type protein and the three engineered mutations. D. Data points and curves for the three engineered mutations (three technical replicates, N = 1), using increased protein concentrations to measure kinetic parameters.

**Extended Data Figure 8.**
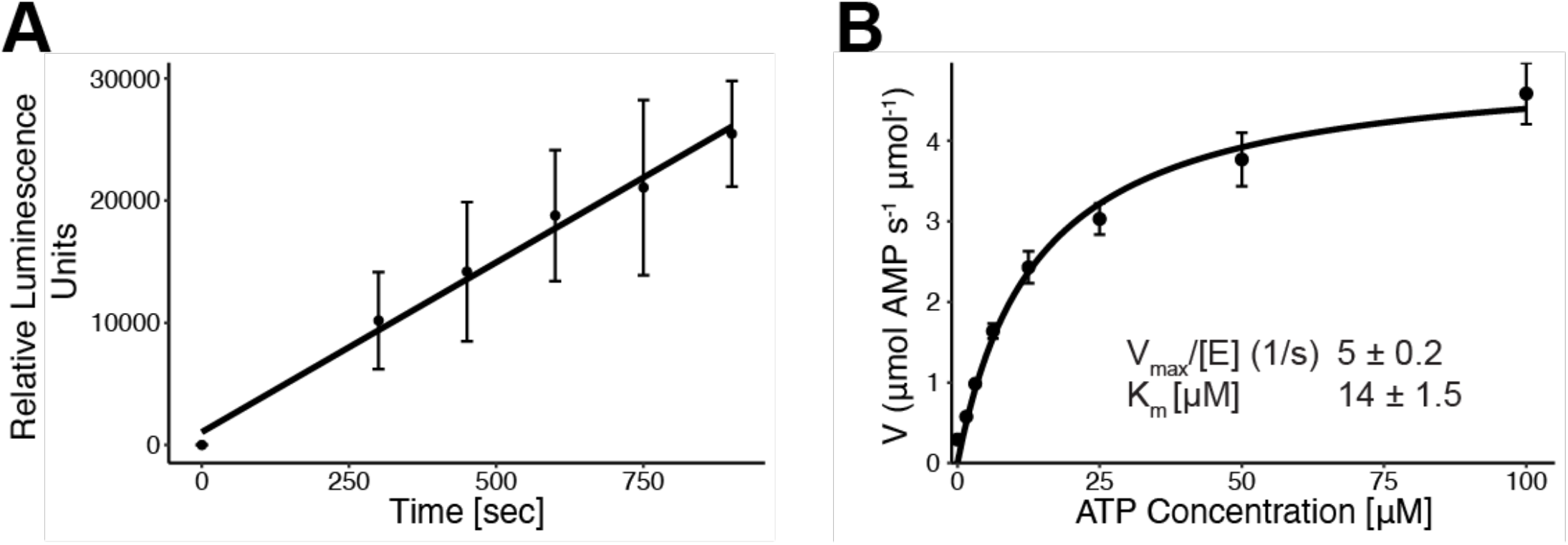
(Figure 4,6) Control assays for catalysis experiments. A. PRPS1 catalysis over time at the lowest ribose-5-phosphate concentration used (100 uM ATP, 1.5 uM ribose-5-phosphate), plotted before conversion to uM AMP. B. PRPS1 kinetic analysis varying ATP concentration and holding ribose-5-phosphate at 100 uM. Calculated kinetic parameters in inset.

**Extended Data Figure 9.**
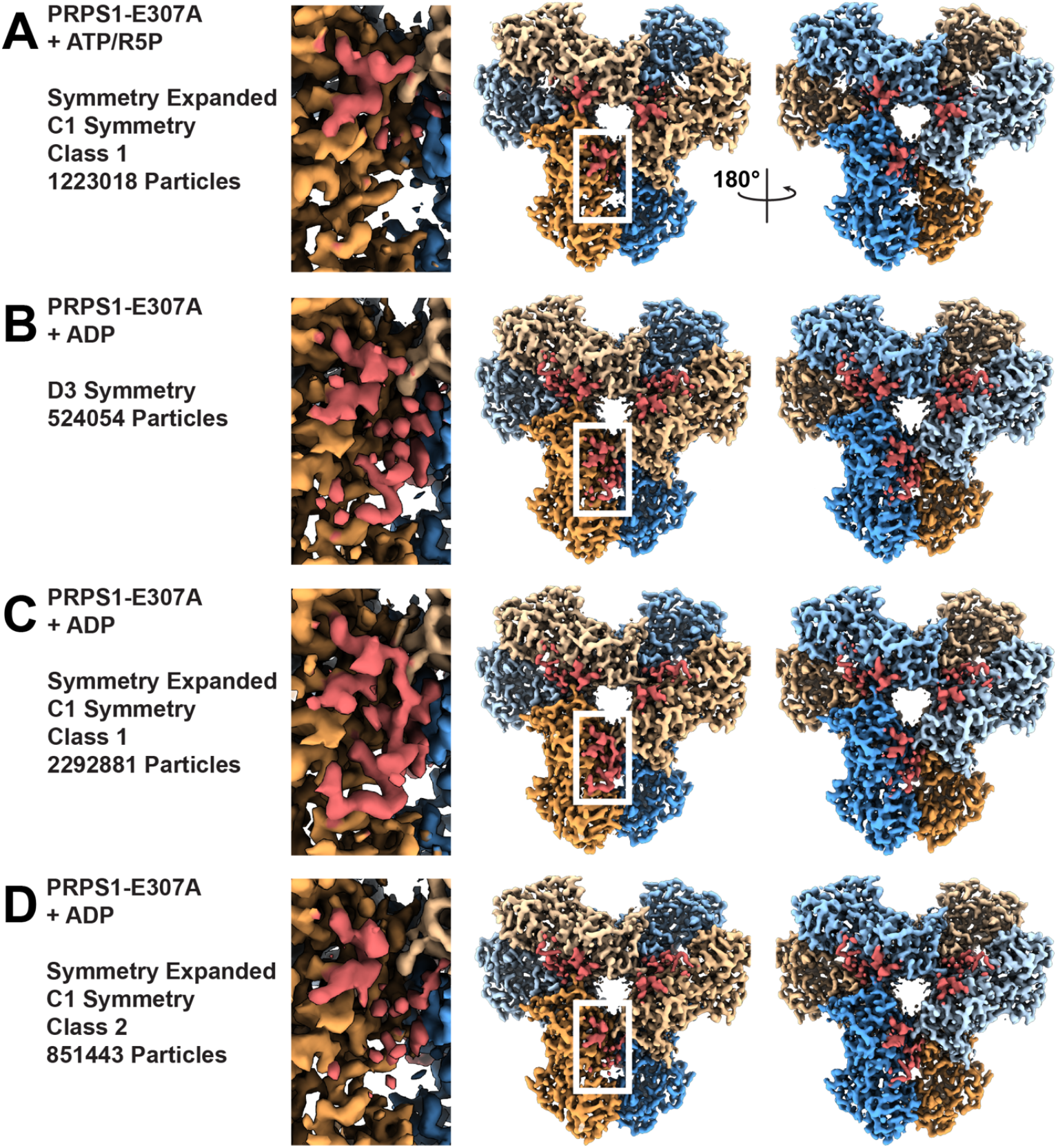
(Figure 5) Volumes for C-termini of PRPS1-E307A mutations. A-D. Panel detailing PRPS1-E307A maps and models. *Left*: Dataset, symmetry, and number of particles included in the map. *Middle*: Insert showing volume of C-termini of protomer a (area highlighted in box on right panel). *Right*: View of both faces of volume from dataset PRPS1-E307A datasets. C-termini are highlighted in red.

**Extended Data Figure 10:**
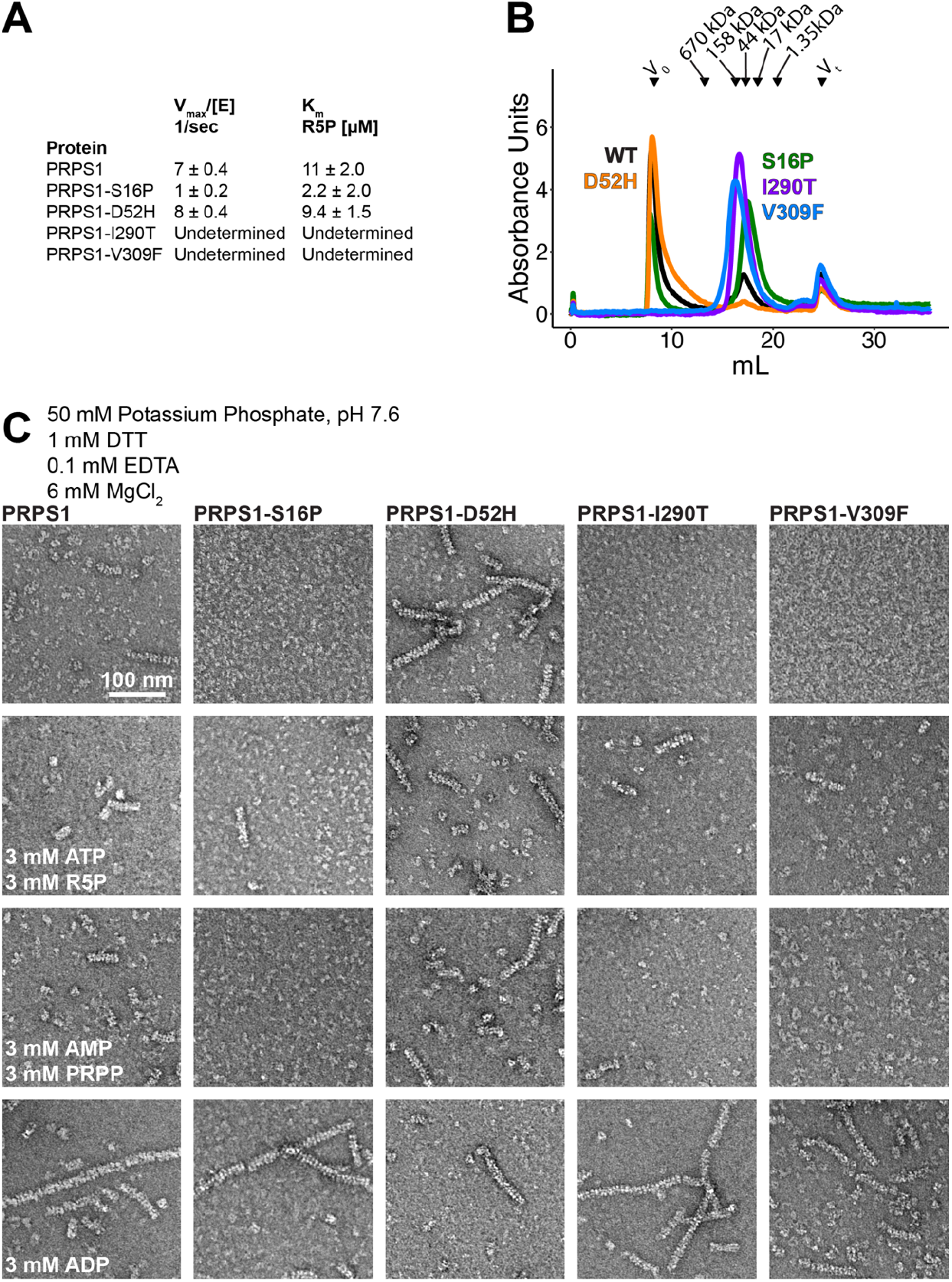
(Figure 6) Filament formation in PRPS1 disease mutants. A. Kinetic parameters for the wild type protein and the four disease mutations as determined from the data shown in Main Text Figure 6. B. Chromatography curves from a Superose 6 of PRPS1 and four disease mutations. C. Panel of negative stain EM of PRPS1 disease mutations in phosphate buffer in the presence of the indicated ligands.

## Supplementary Movies

<u>Supplementary Movie 1: Morph between phosphate-bound PRPS1 filament and ADP-bound PRPS1 filament</u>. Cartoon of backbone atoms, with the protomers colored in orange and blue, and a phosphate positioned in the allosteric site for reference. Filaments have been aligned at the central interface. The movie progresses as follows: phosphate-bound to ADP-bound to phosphate-bound to ADP-bound.

<u>Supplementary Movie 2: Morph between the open and closed conformations of PRPS1 incubated with ATP and ribose-5-phosphate.</u> Top and side views of cartoon of backbone atoms of hexamer, with the protomers colored in orange and blue and the catalytic loops in the a and b protomers in dark orange and dark blue, respectively. Active site ligands are shown in yellow. Hexamers have been aligned on protomers c-f. The movie progresses from protomer b in the open-loop conformation to protomer b in the closed-loop conformation, back to protomer b in the open-loop conformation to protomer b in the closed-loop conformation.

